# Simultaneous measurement of biochemical phenotypes and gene expression in single cells

**DOI:** 10.1101/820233

**Authors:** Amanda L. Richer, Kent A. Riemondy, Lakotah Hardie, Jay R. Hesselberth

## Abstract

Methods to measure heterogeneity among cells are rapidly transforming our understanding of biology but are currently limited to molecular abundance measurements. We developed an approach to simultaneously measure biochemical activities and mRNA abundance in single cells to understand the heterogeneity of DNA repair activities across thousands of human lymphocytes, identifying known and novel cell-type-specific DNA repair phenotypes. Our approach provides a general framework for understanding functional heterogeneity among single cells.

---

New methods to study heterogeneity at cellular resolution measure differences in gene expression ^1–4^, chromatin accessibility ^5^, and protein levels ^6^ across thousands to millions of cells to understand developmental trajectories of tissues, tumors, and whole organisms. But these methods only measure static levels of DNA, RNA, and proteins, limiting our ability to extract dynamic information from individual cells.

We developed a functional assay as a new modality for single-cell experiments. Our key innovation is that, instead of measuring the abundance of molecules—i.e., levels of DNA, RNA, or protein—from single cells and predicting functional states, we directly measure enzymatic activities present in single cells by analyzing the conversion of substrates to intermediates and products in single-cell extracts within a high-throughput DNA sequencing experiment. Our approach is compatible with existing platforms that measure gene expression at single-cell resolution and can measure many different enzymatic activities simultaneously, querying different biochemical activities by combining unique substrates.

We measured DNA repair activities in single cells because the enzymatic substrate (i.e., a DNA lesion to be repaired by cellular enzymes) yields a product that can be directly analyzed by DNA sequencing. DNA damage is repaired by multiple different and often redundant pathways including base excision repair, nucleotide excision repair, mismatch repair, and direct reversal ^7^. Current methods to study DNA repair in cells and cell extracts use synthetic DNA substrates to measure repair activities ^8,9^, but these approaches do not scale to multiple measurements (i.e., gene expression and biochemical activities) from the same cell, and their reliance on substrate transfection precludes facile application to primary cells.

We included synthetic DNA hairpins with polyadenylate tails and DNA lesions at defined positions (**Fig. 1a**) in a single-cell mRNA sequencing experiment and developed a protocol to capture incised DNA repair intermediates and products from single-cell extracts by library preparation and sequencing (**Fig. 1b** and **Supplementary Fig. 1**). Because our method employs massively-parallel measurement of strand incision on DNA hairpins, we named it “Haircut”.

**Figure 1.**
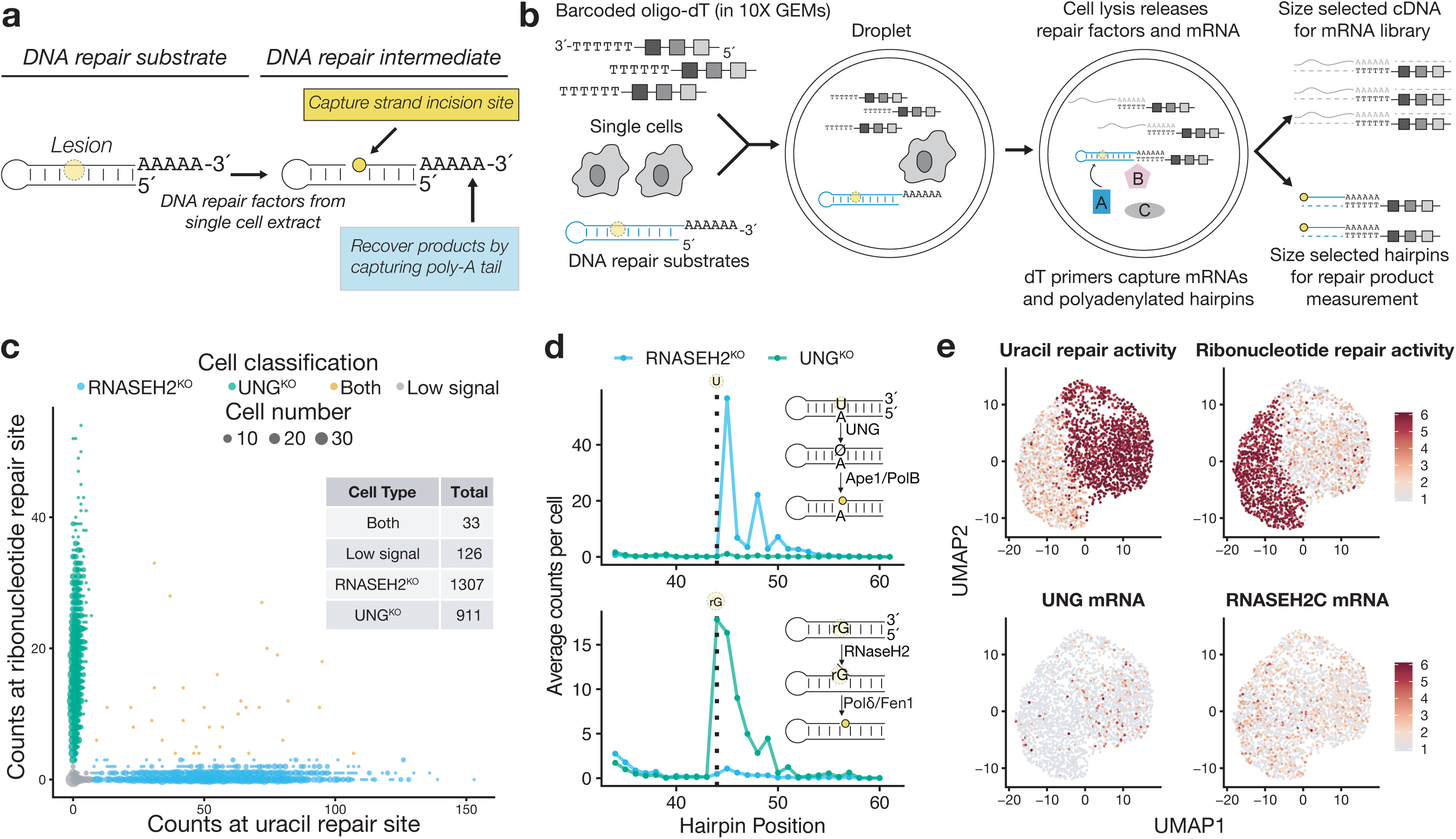
Development and validation of a single-cell assay for measuring DNA repair capacity. **a.** Schematic of DNA repair substrates used in single-cell Haircut. Strand incision generates a new 5′-end whose position is captured by cDNA synthesis with barcoded oligonucleotides. **b.** Overview of single-cell Haircut. After droplet formation, cell lysis creates a ∼50 pL reaction wherein enzymes contributed by the cell catalyze substrate turnover (i.e., strand incision for specific DNA repair substrates). Repair products and mRNAs are converted to cDNA with barcoded oligo-dT primers and separated by size to enable separate library preparations. The cDNA for each fraction is analyzed to identify the cell barcode and either mRNA abundance or the amount of enzymatic activity (i.e., number of strand incisions). **c.** Polyadenylated hairpins containing a single uracil or ribonucleotide were added to a single-cell suspension of a mixture of Hap1 cells containing null alleles of either UNG or RNASEH2C prior to capture in a 10x Genomics 3′ Gene Expression experiment. Sequences from the DNA repair fraction in **(b**) were grouped based on their cell barcodes, and the level of strand incision for uracil and ribonucleotide substrates was used to classify cells as either UNG^KO^ (green) or RNASEH2C^KO^ (blue) based on strand incision activity (UMI counts at position 44 for ribonucleotide, position 45 for uracil) greater than 5% of the maximum for all cells. Cells that fall on the x- or y-axis are single RNASEH2C^KO^ and UNG^KO^ cells, respectively; doublets are in yellow; and cells with low signal (<5% of the max for both activities) are in grey. **d.** Aggregate counts of strand incision events are plotted against hairpin position for cells classified in (**c**) on the U:A substrate (top) and rG:C substrate (bottom). The vertical dashed line notes the position of the uracil and ribonucleotide (position 44 in both cases). UNG^KO^ cells fail to incise uracil-containing hairpins (green line in top panel) and RNASEH2C^KO^ cells fail to incise ribonucleotide-containing hairpins (blue line in bottom panel). The predominant signal at position 45 (top) reflects UNG conversion of the uracil (position 44) to an abasic site, followed by removal of the abasic site by Ape-1 or Pol β. The predominant signal at position 44 (bottom) reflects 5′-incision of the ribonucleotide at position 44 followed by copying of the rG template by reverse transcriptase in the droplet. Additional processing of incised repair intermediates yields signals at positions 3′ of the lesion (signals at positions 46-50 in the uracil substrate (top), and positions 45-50 in the ribonucleotide substrate (bottom)). **e.** Variable mRNA expression from cells classified in (**c**) was used to calculate a UMAP projection (identical coordinates in all 4 panels). Uracil repair activity (natural logarithm of strand incision counts (position 45 for uracil substrate and position 44 for ribonucleotide substrate; **d**) divided by total counts for that cell multiplied by a scaling factor of 10^4^; top left) and ribonucleotide repair activity (top right) were superimposed in a grey-to-red scale and delineate two major cell types in the experiment; 90% of cells in each class have a scored activity. Levels of UNG and RNASEH2C mRNA (natural logarithm of mRNA counts divided by total counts for that cell multiplied by a scaling factor of 10^4^) are plotted on the bottom panels are not sufficient to classify cell types. Stabilization of the null-mutation-containing RNASEH2C mRNA yields uniform RNASEH2C mRNA detection across both cell types.

Cell mixing experiments confirmed we could measure DNA repair activities at cellular resolution with single-cell Haircut. We added DNA repair substrates with a uracil (U:A base-pair) or ribonucleotide (rG:C) to a single-cell suspension of haploid human UNG^KO^ and RNASEH2C^KO^ knockout cells immediately prior to emulsion formation in a droplet-based single-cell mRNA sequencing experiment (10x Genomics 3′ Gene Expression; **Fig. 1b**). After sequential incubations at 37 °C and 53 °C to first promote endogenous enzymatic activity and then reverse transcription, the emulsion was broken and cDNA molecules synthesized from mRNA and repair substrate templates (the “repair fraction”) were separated by size. The mRNA fraction was subjected to the standard protocol to measure gene expression for single cells, whereas the repair fraction (containing DNA substrates, intermediates, and products) was captured in a modified protocol (**Supplementary Fig. 1**). Analysis of the mRNA fraction by DNA sequencing yielded the mRNA identity, a cell-specific barcode, and a unique molecular identifier (UMI ^10^). Similarly, DNA sequencing of the repair fraction yielded the cell barcode, UMI, and a 5′ position derived from cDNA synthesis on either full-length hairpins or incised repair intermediates and products.

We captured thousands of single cells with expected DNA repair defects: RNASEH2C^KO^ cells could incise uracil but not a ribonucleotide repair substrate, and vice versa for UNG^KO^ cells, with a few droplets containing more than one cell from each genotype and therefore both uracil and ribonucleotide repair activities (**Fig. 1c, d**). We also calculated two-dimensional UMAP projections based on variable mRNA expression and colored individual cells by enzymatic activity (UNG or RNASEH2) and mRNA abundance (**Fig. 1e**). DNA repair activity was robustly detected for each cluster (**Fig. 1e**, top row, **Supplementary Fig. 2**) and was sufficient to assign 75% of cells to the correct cell type using 1,500 reads-per-cell (**Supplementary Fig. 2**), similar to read depths required for cell type classification using gene expression ^11^. While UNG (229 cells) and RNASEH2C (1,075 cells) mRNAs were identified in these cells, they were not variably expressed across cell clusters (**Supplementary Fig. 2**). Moreover, our analysis of RNASEH2C mRNA levels in RNASEH2C^KO^ cells found that whereas the mutation yields cells that lack RNASEH2 activity (**Fig. 1e**, top right), it does not cause mRNA decay (**Supplementary Fig. 3**), with similar mRNA isoform abundance detected in both cell types. Altogether, this experiment illustrates the unique and orthogonal information provided by single-cell biochemical assays, which is especially useful in situations where mRNA expression is not predictive or may even be misleading of functional status.

DNA repair activity measurements in single-cell extracts from human peripheral blood mononuclear cells (PBMCs) with separately barcoded uracil and ribonucleotide repair substrates spanning a 50-fold range in concentration showed that measured DNA repair signals change as a function of substrate concentration and time (**Supplementary Fig. 4**), enabling us to choose a single substrate concentration and time point (10 nM substrates at 60 min) for further experiments.

We next used single-cell Haircut to measure the biochemical heterogeneity of DNA repair in PBMCs using five DNA substrates (unmodified, and containing U:A, U:G, rG:C, and abasic:G lesions; **Fig. 2**). We used single-cell mRNA expression to classify cells based on expression of common cell-type-specific markers (e.g., IL7R for CD4+ T cells, CD14 for monocytes; **Fig. 2a**) and then used these classifications to assign cell-type-specific DNA repair activities (**Fig. 2b-f**). The additional steps required to measure DNA repair in thousands of cells impacted the recovery of gene expression signals, typically reducing the number of genes detected by 50% (**Supplementary Fig. 5**); however, this reduction in detection did not preclude classification of cell types by mRNA expression measured in the same cells (**Fig. 2, Supplementary Fig. 6 and 7**).

**Figure 2.**
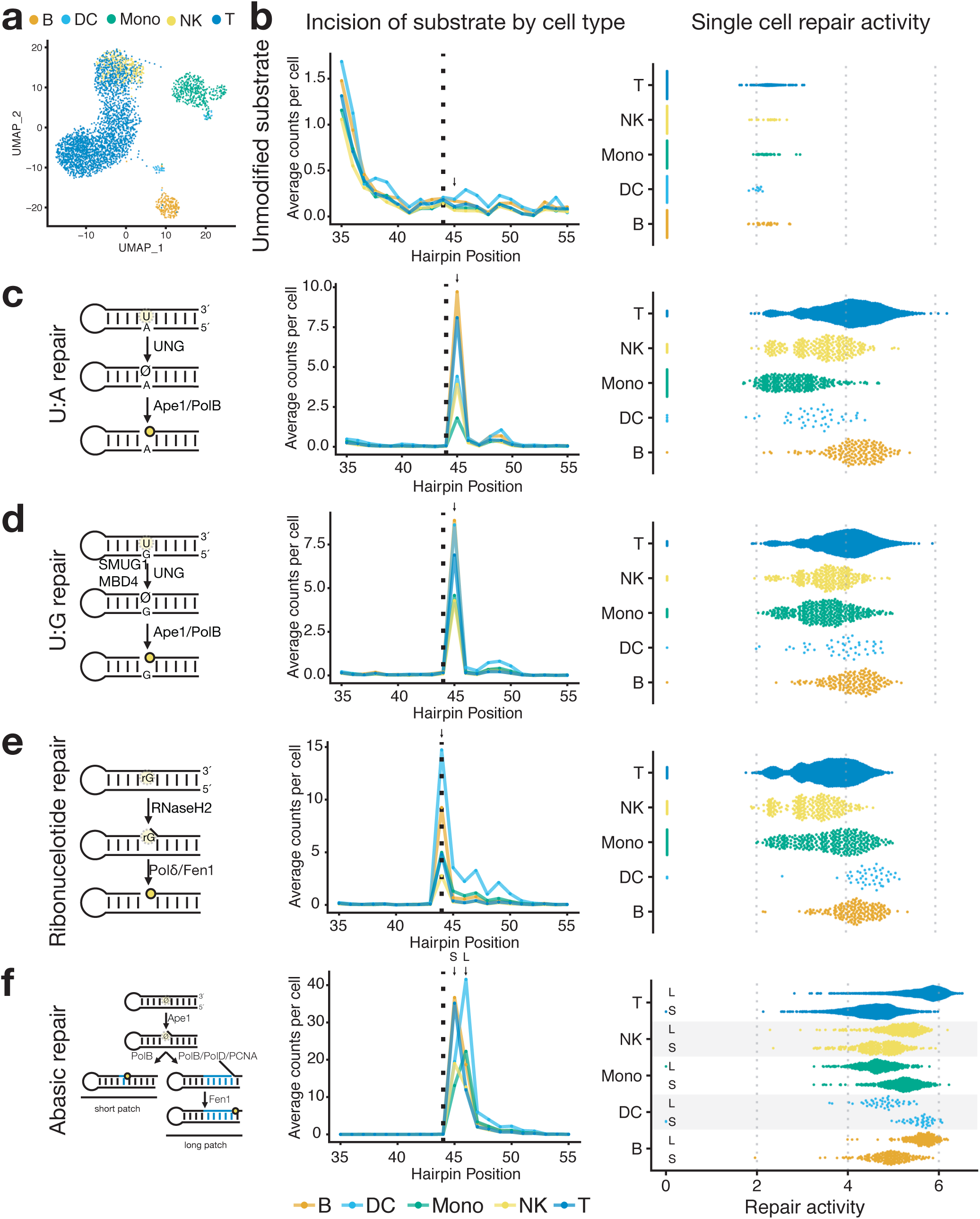
Analysis of DNA repair heterogeneity in human lymphocytes. **a.** Two-dimensional UMAP projection of variable gene expression across 3,298 human PBMCs captured in a 10x Genomics 3′ Gene Expression experiment. Major cell types were determined by marker gene expression (CD19 for B cells; IL7R for T cells; LYZ, FCGR3A, CD14 for monocytes; FCER1A for dendritic cells; GNLY for NK cells). **b.** Single-cell DNA repair activity of an unmodified DNA substrate. The dashed vertical line indicates the position of lesions in other DNA repair substrates (left). Unmodified DNA substrates yield few measured incisions in single-cell Haircut (natural logarithm of strand incision counts (at position 45; arrow, left) divided by total counts for that cell multiplied by a scaling factor of 10^4^), indicating they are not substrates for cellular DNA repair activities. **c.** Repair of a substrates containing a uracil:adenine (U:A) base-pair initiates with UNG-mediated removal of the uracil nucleobase followed by processing of the abasic site by Ape-1 and Pol β (left). Cell-type-specific counts of incision and processing (mean) are plotted against the position of the hairpin and colored as in (**a**) (middle). Single-cell repair activities (natural logarithm of counts at the incision site (arrow) divided by total counts for that cell multiplied by a scaling factor of 10^4^) are plotted for each cell from each cell type (right). Monocytes and dendritic cells have reduced uracil incision relative to other cell types (*P* < 10^−140^ monocytes to T cells, and *P* < 10^−8^ dendritic cells to T cells; Wilcoxon signed-rank test; differences significant across 3 replicates, **Supplementary Table 4**). **d.** Repair of a substrates containing a uracil:guanine base-pair initiates with UNG, SMUG, or MBD4-mediated removal of the uracil nucleobase followed by processing of the abasic site by Ape-1 and Pol β (left). Cell-type-specific incisions are plotted as in (**c**) with a predominant incision site one base downstream of the uracil (arrow, position 45), similar to the U:A substrate (**a**) (middle). The higher uracil repair activity (as defined in **(c)**) on the U:G relative to U:A substrates for monocytes and dendritic cells likely reflects recognition of the U:G substrate by SMUG and MBD4. **e.** Repair of a substrates containing a riboG:C base-pair initiates with RNASEH2-mediated incision 5′ of the ribonucleotide followed by processing by Pol δ and Fen1 (left). Cell-type-specific counts of incision and processing (mean) are plotted against the position of the hairpin and colored as in (**a**) (middle), with the predominant signal at the ribonucleotide, reflecting incision by RNASEH2 and cDNA synthesis using the ribonucleotide template by reverse transcriptase in the droplet (arrow, position 44). B cells and dendritic cells have higher levels of ribonucleotide repair activity (as defined in **(c)**) than other cell types (*P* < 10^−33^ B cells to T cells; *P* < 10^−15^ dendritic cells to T cells; Wilcoxon signed-rank test, differences significant across 3 replicates, **Supplementary Table 4**). **f.** Repair of substrates containing a abasic:guanine base-pair initiates with Ape-1-mediated incision followed by processing of the single-base gap by either Pol β (short-patch repair) or Pol δ/β and Fen1 (long-patch repair) (left). Cell-type-specific incisions are plotted as in (**c**) with a predominant incision sites one or more bases downstream of the abasic, depending on the cell type (middle). For each cell type, the levels of short-patch (top; signals 1 nt downstream of lesion, S arrow, position 45) and long-patch (bottom; signal 2 nt downstream lesion, L arrow, position 46) repair activities (as defined in **(c)**) are plotted (right). Monocytes and dendritic cells have lower levels of short-patch repair relative to other cell types (*P* < 10^−162^ monocytes to T cells; *P* < 10^−19^ dendritic cells to T cells; Wilcoxon signed-rank test, differences significant across 3 replicates, **Supplementary Table 4**).

We found little signal on the unmodified DNA substrate, confirming it is not a repair substrate. In contrast, incision and processing activities measured on uracil (on U:A and U:G substrates), ribonucleotide, and abasic repair substrates matched expected positions based on known repair pathways (left and middle panels in **Fig. 2c-f**) and were only present in cell-associated droplets (as determined from mRNA signals; **Supplementary Fig. 8**). These data revealed unique signatures of DNA repair activities in specific cell types. Monocytes and dendritic cells had low incision activity on the U:A substrate (**Fig. 2c**), resonating with the low level of uracil base excision in monocytes ^12^ and confirming that myeloid lineages have unique uracil repair phenotypes ^13^. However, these differences in uracil repair were not apparent for the U:G substrate, presumably due to redundant activities of SMUG1 ^14^ and MBD4 ^15^ in incising U:G-containing substrates. Dendritic cells demonstrated a unique repair phenotype, with increased levels of DNA substrate processing, measured as increased signals downstream of the position of the synthetic lesion. To explain these differences in DNA repair phenotypes, we examined cell type-specific mRNA expression and found that expression of APEX1, encoding the abasic endonuclease Ape-1, was consistently and uniquely elevated in dendritic cells (**Supplementary Tables 1 and 2**), possibly explaining the increased processing of DNA repair intermediates on U:A, U:G, and abasic substrates in dendritic cells.

B cells and dendritic cells also had higher levels of ribonucleotide repair activity than other cell types (**Fig. 2** and **Supplementary Fig. 6**), and the increase in ribonucleotide repair in B cells and dendritic cells was corroborated by elevated RNASEH2 expression in our (**Supplementary Table 1 and 2**) and previous single-cell mRNA sequencing data sets (**Supplementary Table 3**) ^4^. Increased RNASEH2 activity in B cells may aid processing of R-loops that form during class switching ^16^.

Finally, we focused on cell-type-specific differences in repair of a DNA hairpin containing a synthetic abasic site. These substrates undergo two unique events in droplets: on intact substrates, reverse transcription halts at the abasic site, yielding extension products that map one base downstream of the abasic site (**Fig. 2f**). Alternatively, incision and removal of the abasic site by Pol β and Fen1 ^17^ yields repair intermediates with 5′-ends that map further downstream of the abasic site. Differences in signals from the abasic substrate specifically in monocytes and dendritic cells again indicate that they are more proficient at processing abasic lesions, evidenced by an increase in levels of intermediates 2 nt downstream of the abasic site (position 46; **Fig. 2f** and **Supplementary Fig. 6**), likely due to elevated Ape-1 expression. The unique DNA repair phenotype provided some power for cell type classification (**Supplementary Fig. 7g**). As additional activities are multiplexed with DNA repair, we expect the power of single-cell biochemical measurements for cell type classification will increase.

Our approach to measure heterogeneity of single-cell biochemical phenotypes can be expanded to measure other types of DNA repair activities (e.g., nucleotide excision repair and mismatch repair) and adapted to measure other enzyme classes using substrate-DNA conjugates ^18^, enabling simultaneous measurement of many biochemical activities (e.g., kinases, phosphatases, and proteases) with gene expression at single-cell resolution.

## DATA AVAILABILITY

Sequencing data have been deposited at NCBI GEO under accession GSE130117. A full protocol is available at https://dx.doi.org/10.17504/protocols.io.uhyet7w. An analytical and reproducible pipeline is available at https://github.com/hesselberthlab/sc-haircut.

## ACKNOWLEDGEMENTS

We thank members of the Hesselberth lab for technical discussions, and we thank A. Johnson and S. Ramachandran for comments on the manuscript. This work was supported in part by a National Science Foundation Graduate Research Fellowship (DGE-1553798 to A. R.), the National Institutes of Health (R35 GM119550 to J.H.), the Golfers Against Cancer (Denver Chapter), and the Linda Crnic Institute for Down Syndrome.

## SUPPLEMENTAL METHODS

### Single cell experiments

#### DNA repair substrates for single cell experiments

Oligonucleotides were purchased from IDT (**Supplementary Table 5**). Substrates contain a 5′ and 3′ C3 spacer to prevent exonuclease degradation and reverse transcriptase extension of the substrates. Hairpins were gel purified prior to use in single cell experiments. Briefly, 2-5 nmoles of hairpins were loaded in denaturing buffer (47.5% formamide, 0.05% Orange G) on 8% 19:1 acrylamide (BioRad) TBE-Urea gels (7 M urea, 0.1 M Tris base, 0.1 M boric acid, 2 mM EDTA). Hairpins were visualized with UV shadowing on a TLC Silica gel 60 F₂₅₄ plate (Millipore), cut from the gel, crushed in a 1.5 ml Eppendorf tube and eluted in 400 µl 0.3 M sodium acetate overnight at 37 °C shaking at 400 RPM. Acrylamide was removed using 0.45 μm cellulose acetate filters (Costar). Hairpins were then purified via ethanol precipitation and resuspended in water. The concentration of purified hairpins was determined via absorbance at 260 nm on a Nanodrop 2000 (Thermo Scientific).

#### Preparation of single cell suspensions

Single cell suspensions from cell lines were prepared according to 10x Genomics guidelines. Briefly, cells were quickly washed with 0.25% trypsin (ThermoFisher) and then incubated in 0.25% trypsin for 5 minutes at 37 °C. Trypsin digestion was quenched by the addition of cell culture medium. Cells were isolated by centrifugation at 150 xG for 3 minutes (these same conditions were used for all cell washes). For cell mixing experiments, approximately 10^6^ cells from each knockout cell line (UNG^KO^ or RNASEH2C^KO^) were filtered through a 30 µm strainer and mixed in the same tube. Cells were washed twice with cold PBS containing 0.04% BSA. Cells were resuspended in 500 µl PBS with 0.04% BSA and filtered through a Flowmi™ Tip Strainer. Cells were stained with trypan blue and counted on a hemocytometer. Cell concentration ranged from 400 - 1000 cells per µl and viability was between 80-95%.

Fresh peripheral blood mononuclear cells (PBMC) were isolated from whole blood donated by healthy human donors according to University of Colorado IRB guidelines in sodium heparin tubes. Approximately 5-10 ml of whole blood was diluted with PBS to a total volume of 35 ml. Diluted whole blood was layered over 10 ml Ficoll-Paque PLUS (GE) and centrifuged at 740 xG for 20 minutes with no deceleration. Cells located above the Ficoll layer were removed and washed twice with PBS. Cells were counted and approximately 2 million cells were washed an additional two times with PBS plus 0.04% BSA. Cells were resuspended in 500 µl PBS plus % BSA and run through a Flowmi™ Tip Strainer. Cells were counted on a hemocytometer: cell concentration ranged from 400-1000 cells per µl and viability was between 80-95%.

#### Single cell repair measurements using the 10x Genomics platform

The most current version of this protocol is available at: https://dx.doi.org/10.17504/protocols.io.uhyet7w.

Cells were loaded onto the 10x Genomics single cell 3′ expression kit V2 according to the manufacturer’s instructions (CG 000075 Rev C) with the following exceptions:

1. When preparing the single cell master mix, 5 µl was subtracted from the appropriate volume of nuclease-free water. After the nuclease-free water was added to the master mix and prior to the addition of the single cell suspension, 5 µl of DNA repair substrates were added (see **Supplementary Table 6** for substrate concentrations for each experiment).
2. The GEM-RT incubation was changed to the following:
  2.1. Lid temperature: 53 °C
  2.2. 37 °C for 60 minutes (unless otherwise noted in experiment **Supplementary Fig. 4**)
  2.3. 53 °C for 45 minutes
  2.4. 4 °C Hold and proceed directly to GEM-RT cleanup
3. After GEM-RT cleanup, DNA repair substrates and products were separated from mRNA prior to cDNA amplification. 0.6x volume of AmpureXP was added to the eluted RT products (21 µl AmpureXP to 35 µl RT product) and incubated for 5 minutes at room temperature. The sample was placed on a magnetic strip (High on 10x Magnetic Separator) until the liquid was clear. The supernatant was transferred to a new tube since it contained the DNA repair substrates and products. The beads containing the RT products were washed twice with 150 µl of 80% ethanol, then dried for 2 minutes at room temperature, and eluted in 35.5 µl of Elution Solution 1 according to 10x SPRIselect cleanup protocols. This fraction was used to prepare the mRNA expression library according to manufacturer’s instructions. The supernatant was cleaned up by added 1.8x of the original volume of AmpureXP (42 µl), mixed, and then incubated for 5 minutes at room temperature. The sample was placed on a magnetic strip until the liquid was clear. The supernatant was discarded and the beads were washed twice with 150 µl 80% ethanol. The beads were dried at room temperature for 2 minutes, then eluted in 20 µl water. This fraction was used to prepare the DNA repair libraries.

#### Preparation of DNA repair libraries from single cells

The DNA repair libraries were prepared with the following steps:

1. *End repair*: 20 µl of the purified DNA repair libraries were added to an end repair reaction with a total volume of 30 µl (NEBNext End repair Module E6050) and incubated for 30 minutes at 20° C.
2. *Clean up by precipitation*: 120 µl of 0.3 M sodium acetate and 400 µl 100% ethanol were added to the end repair reaction (step 1). The reaction was allowed to precipitate at −20 °C for at least 30 minutes. Samples were centrifuged at >10000 XG for 10 minutes and the supernatant was removed and the pellet was washed with 500 µl of 70% ethanol and centrifuged at >10000 xG for 10 minutes. The supernatant was removed and the pellet was dried for 2 minutes at room temperature. The pellet was resuspended in 20 µl nuclease-free water.
3. *A-tailing:* 15 µl of the end repaired DNA repair library was added to an A-tailing reaction with a total volume of 20 µl (1x Blue Buffer (Enzymatics), 400 µM dATP, 5 units Klenow 3′-5′ exo- (Enzymatics)) for 30 minutes at 37 °C. The A-tailing reaction was cleaned up using precipitation as in step 2.
4. *Adapter ligation*: 13 µl of the A-tailing reaction was added to an Illumina Y adapter ligation reaction with a total volume of 20 µl (66 mM Tris-HCl, 10 mM MgCl_2_, 1 mM DTT, 1 mM ATP, 7.5% PEG 6000, pH 7.6, 0.3 µM annealed Y adapters, 600 units Rapid T4 DNA Ligase (Enzymatics)) and incubated at 25 °C for 30 minutes. The ligation reaction was purified using 1.8 x volume of Agencourt AMPure XP (Beckman Coulter) beads as described by the manufacturer. The reaction was eluted in 20 µl nuclease-free water.
5. *Illumina TruSeq PCR*: 13 µl of the purified ligation reaction was added to a PCR reaction with a total volume of 50 µl (1x Phusion HF buffer (NEB), 200 µM dNTPs, 0.6 µM ILMN PCR primers (F and R), 2 units Phusion High Fidelity DNA polymerase). 14-20 cycles of PCR were done with 98 °C melting temperature for 15 seconds, 65 °C annealing temperature for 15 seconds, and 72 °C extension temperature for 15 seconds.
6. *PCR cleanup and sequencing:* The DNA repair library was purified using 1.2x volume Agencourt AMPure XP (Beckman Coulter) beads as described by the manufacturer. The DNA repair library was quantified using the Qubit HS dsDNA fluorometric quantitation kit (Thermo Scientific). 1 µl of the DNA repair library was analyzed on the Agilent D1000 Tapestation. The DNA repair library was ∼230-250 base pairs. The DNA repair library was paired end sequenced on a NovaSeq 6000 system with 2×150 base pair read lengths at the University of Colorado Anschutz Medical Campus Genomics and Microarray core. Each library was sequenced with at least 10 million reads per sample.

#### Single cell data processing

Data processing scripts are available at https://github.com/hesselberthlab/sc-haircut. Briefly, FASTQ files from the 10x mRNA libraries were processed using the cellranger count pipeline (v3.0.2). Reads were aligned to the GRCh38 reference. For the repair libraries, the cell barcodes and UMIs were extracted from R1 using umi_tools ^19^. All of the known 10x cell barcodes were provided as the whitelist. R2 was trimmed to remove the 3′ polyA sequence and the 5′ template switching sequence. Then R2 was aligned to a hairpin reference fasta file using bowtie2 (v2.3.2) ^20^, no reverse complement alignment was allowed to ensure sequences aligned in the correct orientation to the reference. The chromosome (same as substrate name) and 5′ alignment position were concatenated and added to the bam file in the XT flag. UMIs were grouped and appended to the BAM files as a tag using umi_tools group. UMIs were counted per cell per hairpin position using umi_tools count. The table output was converted into a sparse matrix and filtered by matching cell barcodes found in the cellranger filtered feature matrix output using functions in the scrunchy R package (https://github.com/hesselberthlab/scrunchy).

##### Seurat

Downstream analysis of RNA and repair data was performed using the Seurat R package (v3.0.0) ^21^. Raw, filtered counts for repair was added to the same Seurat object as gene expression. Gene expression counts and repair counts were log normalized (LogNormalize) where feature counts for each cell are divided by the total counts for that cell and multiplied by a scaling factor (10^4^) and then natural-log transformed. PBMC samples were filtered for number of genes per cell >150-200 and <2000-2500 and for percent mitochondrial reads <15%-25%. Gene expression data was scaled and centered (ScaleData). 5000 variable features (FindVariableFeatures) were used for PCA calculation (RunPCA) and the first 10-20 principal components were used to find clusters (FindNeighbors, FindClusters) and calculate uniform manifold approximation and projection (UMAP) (RunUMAP). Cell types were identified using the Seurat functions FindTransferAnchors and TransferData ^21^. Reference PBMC data was downloaded from Seurat vignette and used as reference for PBMC cell types (https://satijalab.org/seurat/v3.1/pbmc3k_tutorial.html; https://support.10xgenomics.com/single-cell-gene-expression/datasets/1.1.0/pbmc3k). Cells were filtered to exclude platelets unless otherwise noted (**Supplementary Fig. 7**). Significant differences and fold changes in repair activities and gene expression between cell types were calculated using Wilcoxon Rank Sum test (FindMarkers, FindAllMarkers) for all pairwise combinations (**Supplementary Table 1, 2, and 4**). In the cell mixing experiment (**Fig. 1**), cell types were determined by repair activities. Knockout cells were identified if counts at the repair site (position 44 for ribonucleotide and position 45 for uracil) for one repair activity was > 5% of the maximum for the repair activity and the other was < 5% of the maximum. If both repair activity counts were >5% of the maximum, that cell was considered a doublet and if both repair activity counts were <5% of the maximum that cell was not classified. Filtered single cell gene expression matrices from previously published data ^4^ for **Supplementary Table 3** were downloaded from 10x Genomics (https://support.10xgenomics.com/single-cell-gene-expression/datasets/3.0.2/5k_pbmc_v3_nextgem, https://support.10xgenomics.com/single-cell-gene-expression/datasets/3.1.0/5k_pbmc_protein_v3, https://support.10xgenomics.com/single-cell-gene-expression/datasets/3.0.0/pbmc_10k_v3) and analyzed the same as above.

##### Genome coverage

To calculate genome coverage for cell mixing experiment (**Supplementary Fig. 3**), BAM files produced by cellranger were split into cell type (as assigned above and in **Fig. 1**) specific BAM files by cell barcodes using samtools view (v1.9) ^22^. Bulk genome coverage was calculated for UNG^KO^ or RNASEH2C^KO^ cells using bedtools genomecov (v2.26.0) ^23^. Coverage was visualized with the UCSC Genome Browser ^24^.

##### Quality control metrics

Quality control metrics (e.g. number of genes, median genes per cell, median UMIs per cell, etc) were calculated in the cellranger pipeline (metrics_summary.csv). To compare recovery of mRNA with and without the addition of DNA repair substrates, we used PBMC reference data from 10x Genomics with roughly the same number of cells (https://support.10xgenomics.com/single-cell-gene-expression/datasets/2.1.0/pbmc4k; https://support.10xgenomics.com/single-cell-gene-expression/datasets/3.0.0/pbmc_1k_v2). FASTQ files downloaded from 10x data were subsampled to the same read depth as our samples then analyzed using cellranger.

##### Cell type classification with repair data

To determine whether DNA repair activities are useful in classifying PBMC cell types, we used the PBMC replicates 1 and 2 (**Fig. 2** and **Supplementary Fig. 6**, left). True cell types were determined using reference PBMC data from 10x Genomics as described in Seurat analysis section above. All other cell type classifications were compared to these reference cell types. Next, cell types were determined by renaming the defined Seurat clusters as the majority cell type present within each cluster (**Supplementary Fig. 7b**). Additionally, mRNA (**Supplementary Fig. 7c**) and/or repair data (**Supplementary Fig. 7d,e**) from PBMC replicate 1 was used as the reference input for assigning cell types to PBMC replicate 2 using Seurat’s FindTransferAnchors and TransferData functions. Cell types were randomly reassigned using R (sample) (**Supplementary Fig. 7f**). True positive, true negative, false positive, and false negative numbers were calculated for each cell type for each classification method. These numbers were then used to calculate true positive rate and false positive rate for each cell type for each classification method (**Supplementary Fig. 7g**).

##### Empty drops vs cells analysis

To determine biological vs background activity in single cells, we calculated hairpin coverage in empty drops (**Supplementary Fig. 8**). Empty drops were determined by filtering out cell-associated barcodes from the repair matrix. The resulting repair matrix contains many barcodes that are associated with only a single UMI, so the matrix was filtered by descending UMI counts to the same number as cell-associated barcodes. This repair matrix from empty drops was used as the input to calculate empty drop signal across the hairpin by calculating the mean count across all drops at each hairpin position.

##### Haircut signal detection sensitivity

To determine the lower limit of haircut signal detection suitable for classifying cells as either UNG^KO^ or RNASEH2C^KO^ read alignments were randomly downsampled using samtools view (v1.9) ^22^. The downsampled BAM files were then processed using the haircut single cell processing pipeline to produce haircut signal matrices. Cells were classified as either UNG^KO^, RNASEH2C^KO^, doublets, or low signal using the same cutoffs described in the Seurat analysis methods.

##### Direct reversal substrate and 5′ biotin cleanup

To measure repair of direct reversal substrates we included O^6^methyl-G substrate in PBMC experiments (**Supplementary Fig. 6 and 8**). Prior to end repair, the repair fraction was digested with PstI (NEB, 20 U in 1x Cutsmart buffer) at 37 °C for 60 minutes. The reaction was cleaned up by precipitation, followed by the above steps starting at end repair. To remove background signal on the 5′ end of the substrates, we included a uracil substrate with a 5′ biotin in PBMC experiments (**Supplementary Fig. 6 and 8**). To remove uncleaved substrate, prior to end repair, the repair fraction was incubated with Dynabeads™ M-270 Streptavidin (5 μg, Thermo) for 5 minutes at room temperature. Following incubation, the beads were discarded and the supernatant was cleaned with precipitation. The remainder of the protocol proceeded starting from end repair.

#### Oligonucleotides for repair libraries

Other oligonucleotides used in the library preparation can be found in **Supplementary Table 5.** To anneal Y adapters, 100 µM Y adapter 1 and 2 were mixed in equimolar concentration in 10 mM Tris-HCl pH 7.5, 50 mM NaCl and heated to 95 °C and cooled to 4 °C over 5 minutes. Annealed adapters were diluted to 10 µM final concentration in 10 mM Tris-HCl pH 7.5, 50 mM NaCl. Annealed adapters were stored at −20 °C until use.

#### Cell lines and cell culture

Hap1 UNG^KO^ (HZGHC001531c012) and RNASEH2C^KO^ (HZGHC004633c003) cells were purchased from Horizon Discovery. Cell lines were cultured in IMDM (Gibco, purchased from ThermoFisher) supplemented with 10% FBS (ThermoFisher) and Penicillin-Streptomycin (ThermoFisher) at 37 °C with 5% CO_2_.

#### RT-qPCR

Total RNA from cells was isolated using TRIzol reagent (ThermoFisher) according to manufacturer’s instructions. Total RNA (5 µg) was treated with TURBO DNAse (2 U) (ThermoFisher) according to manufacturer’s instructions. Following DNAse treatment, 1 µg of total RNA was reverse transcribed using Superscript II (200 U, ThermoFisher) and random hexamers primers (0.5 µM, ThermoFisher) to make cDNA. The cDNA was then used for quantitative PCR (qPCR) using Sso Advanced Universal SYBR Green Supermix (Bio Rad) and cycled on a Bio Rad C1000 384-well thermal cycler and plate reader. qPCR experiments were done in technical triplicate and biological duplicates.

## SUPPLEMENTAL TABLES

### TABLE 1. Differential gene expression

Significant differences and fold changes in gene expression between cell types and all other cells were calculated using Wilcoxon Rank Sum test (FindAllMarkers, Seurat v3.0.0 ^21^) for all cell types. Statistics were calculated for each replicate individually (sample: pbmc1, pbmc2, pbmc3) and for all samples combined (sample: all cells). Column information is as follows (from Seurat documentation for FindMarkers):

- gene: Gene name
- cluster: Cell type determined by mRNA expression
- p_val: p value of Wilcoxon Rank Sum test
- avg_logFC: The log fold-change of average expression between group and all other cells. Positive values indicate that the gene is more highly expressed in the first group.
- pct.1: Percentage of cells where the gene was detected in the group.
- pct.2: Percentage of all other cells where the gene was detected.
- p_val_adj: Adjusted p-value, based on bonferroni correction using all genes in the dataset
- sample: The replicate for which the statistics were calculated. “All_cells” is a combination of all replicates.

### TABLE 2. Differential gene expression for base excision repair genes

Significant differences and fold changes in gene expression between cell types and all other cells were calculated using Wilcoxon Rank Sum test (using the FindAllMarkers function from Seurat v3.0.0 ^21^) for all cell types. The result was filtered for genes within the KEGG base excision repair pathway (hsa03410). Statistics were calculated for each replicate individually (sample: pbmc1, pbmc2, pbmc3) and for all samples combined (sample: all cells). Columns are the same as in **Supplementary Table 1**.

### TABLE 3. Differential gene expression for base excision repair genes from reference single cell mRNA datasets

Significant differences and fold changes in gene expression between cell types and all other cells and all pairwise comparisons were calculated using Wilcoxon Rank Sum test (FindAllMarkers, FindMarkers, Seurat v3.0.0 ^21^) for all cell types. Samples were downloaded from 10x Genomics ^4^. Data from Donors A, B and C (frozen) ^4^ were used as well as reference data from 5K PBMC using NextGem technology (https://support.10xgenomics.com/single-cell-gene-expression/datasets/3.0.2/5k_pbmc_v3_nextgem), 5K PBMC using V3 chemistry (https://support.10xgenomics.com/single-cell-gene-expression/datasets/3.1.0/5k_pbmc_protein_v3), and 10K PBMC using V3 chemistry (https://support.10xgenomics.com/single-cell-gene-expression/datasets/3.0.0/pbmc_10k_v3). Results were filtered for adjusted *P* < 0.05 and genes within the KEGG base excision repair pathway (hsa03410). Columns are the same as Table 1 with the addition of celltype1 and celltype2 indicating the pairwise comparisons.

### TABLE 4. Differential repair calculated for biological repair positions

Significant differences and fold changes in repair activities between cell types and all other cells and all pairwise comparisons were calculated using Wilcoxon Rank Sum test (FindAllMarkers, FindMarkers, Seurat v3.0.0 ^21^) for all cell types. Statistics were calculated for each replicate individually (sample: pbmc1, pbmc2, pbmc3) and for all samples combined (sample: all_cells). The result was filtered for adjusted *P* < 0.05 and repair positions (**Fig. 2**). Column information is the same as **Supplementary Table 1**. With the exception of the repair column which is a concatenation of repair substrate and position and the addition of celltype1 and celltype2 to indicate the pairwise comparisons.

### TABLE 5. Oligonucleotides for single cell DNA repair measurements

Substrates and other oligonucleotides used in single-cell Haircut experiments. We did not measure biological activity from the following substrates in single cells: C:I, T:I, T:ethenoA, C:O6mG, A:hmU (**Supplementary Fig. 8**). These substrates were included in single-cell Haircut experiments with human PBMCs (**Supplementary Table 6**).

### TABLE 6. DNA repair substrate experimental conditions

This table contains the substrates and concentrations used in single-cell Haircut experiments.

## SUPPLEMENTARY FIGURE LEGENDS

**Supplementary Figure 1.**
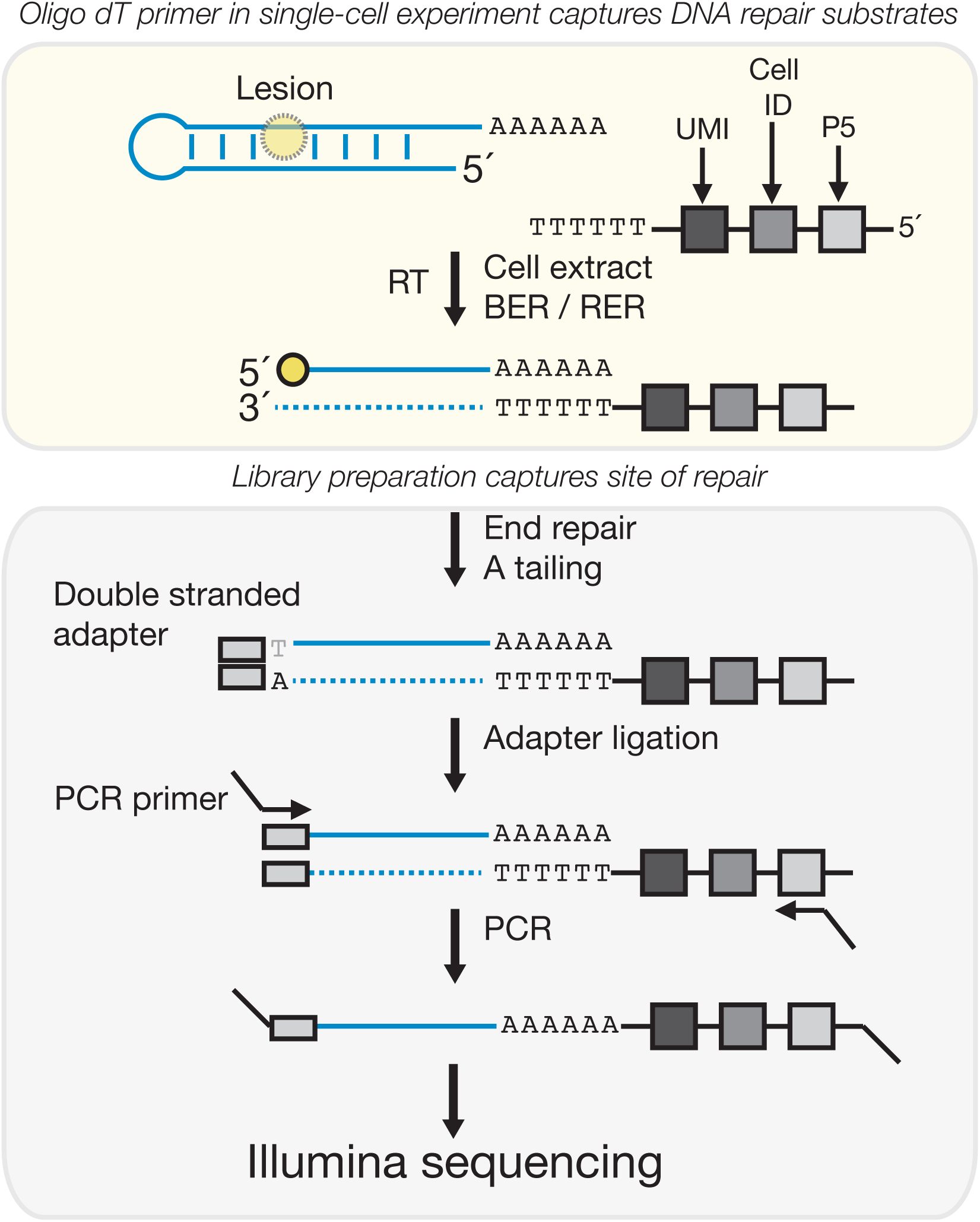
Schematic of single-cell Haircut library preparation. DNA repair substrates are added to a 10x Genomics Chromium Single Cell 3′ v2 kit. Within each droplet, DNA repair substrates are exposed to cell extract containing active DNA repair enzymes, a reverse transcriptase, and an oligo-dT reverse transcription primer. DNA repair enzymes initiate DNA repair on the substrates creating a strand incision, oligo-dT primer and reverse transcriptase capture the DNA repair intermediates along with cellular mRNAs. After the emulsion is broken, DNA repair substrates are separated from mRNAs by size. The 5′ site of strand incision is captured through end repair, A-tailing, ligation of a TruSeq adapter, followed by PCR. This library is compatible with next generation sequencing.

**Supplementary Figure 2.**
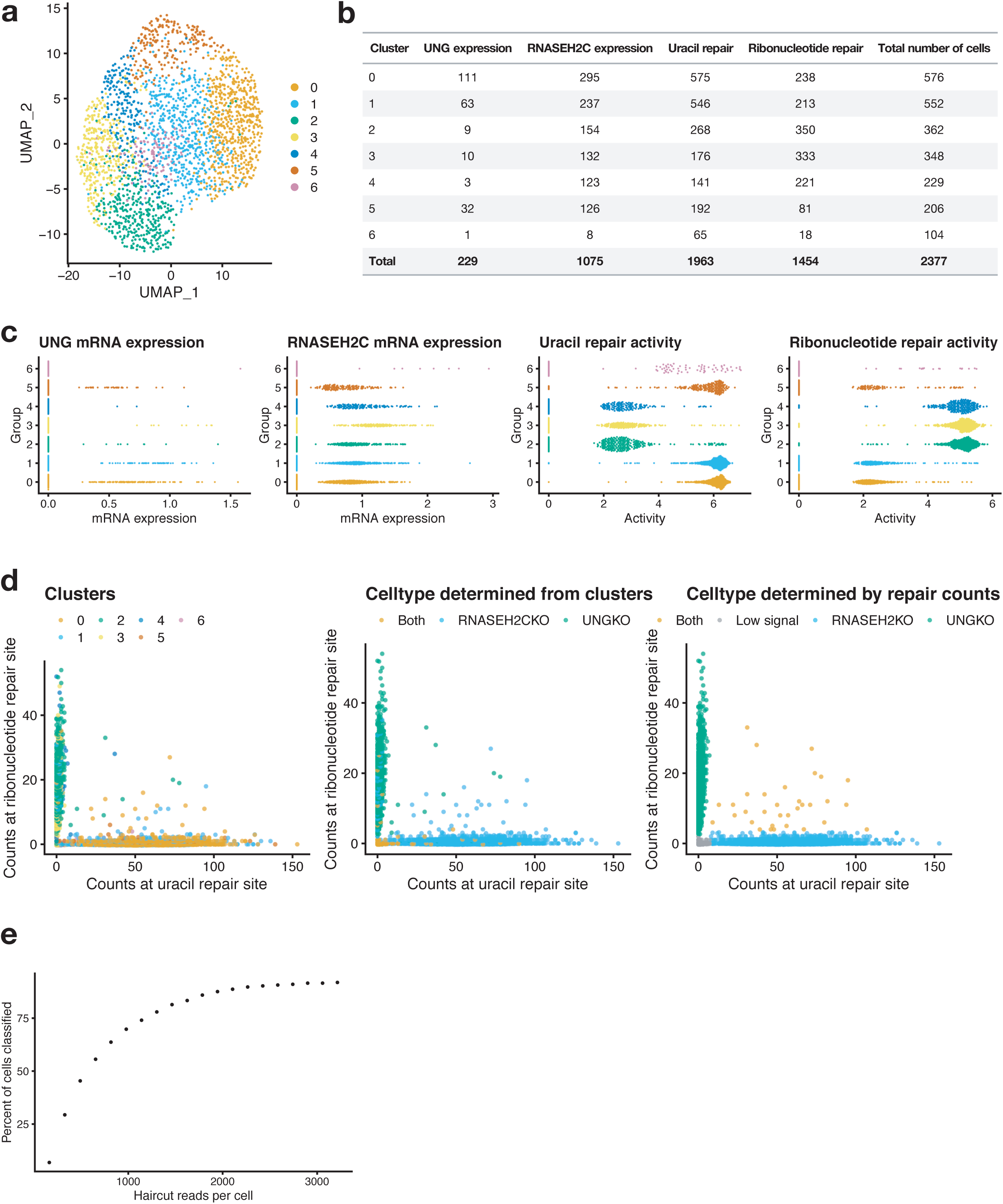
DNA repair measurements determine cell types in a cell mixing experiment. **a.** UMAP plot of cell mixing experiment (**Fig. 1**). mRNA expression was used to calculate UMAP projections and cluster cells. Cells were clustered using an unsupervised shared nearest neighbors method in Seurat (FindNeighbors, FindClusters). Cells were colored by the resulting cluster numbers. **b.** A table of cells expressing UNG mRNA, RNASEH2C mRNA, uracil repair activity, or ribonucleotide repair activity in each cluster. UNG mRNA was detected in < 10% of cells while RNASEH2C mRNA was detected in ∼45% of cells. One or both repair activities were measured in >90% of cells. **c.** Beeswarm plots of UNG mRNA expression, RNASEH2C mRNA expression, uracil repair activity, or ribonucleotide repair activity by cluster. To determine if UNG and RNASEH2C mRNA expression could be used to assign cell types, we made beeswarm plots of UNG expression and RNASEH2C expression and attempted to assign cell types based on expression levels by cluster (left 2 plots). Due to few cells with UNG or RNASEH2C mRNA measurements we could not assign cell types independent of repair measurements. To assign cell types by repair activity, we made beeswarm plots of repair activities by clusters (right 2 plots). Clusters with high uracil repair activity and low ribonucleotide repair activity were assigned as RNASEH2C^KO^ cells (clusters 0, 1, and 5). Clusters with high ribonucleotide repair activity and low uracil repair activity were assigned as UNG^KO^ (clusters 2, 3, and 4). Cluster 6 was was assigned as having both repair activities. **d.** Counts at ribonucleotide repair site and uracil repair site were plotted and colored by cluster (left), cell type as determined above (**c**) (middle), and cell types as determined in **Fig. 1** (right). Cell types as determined in **Fig. 1** show the greatest segregation in these plots. **e.** BAM files from the cell mixing experiment in **Fig. 1** were down sampled to different sequencing depths to determine how many reads per cell are required for cell type classification. At each threshold, cell types were classified as in **Fig. 1** and the percent of cells that can be classified as either UNG^KO^ or RNASEH2C^KO^ is plotted as a function of sequencing depth. Approximately 75% of the 2,377 cells are correctly classified at 1,500 reads per cell.

**Supplementary Figure 3.**
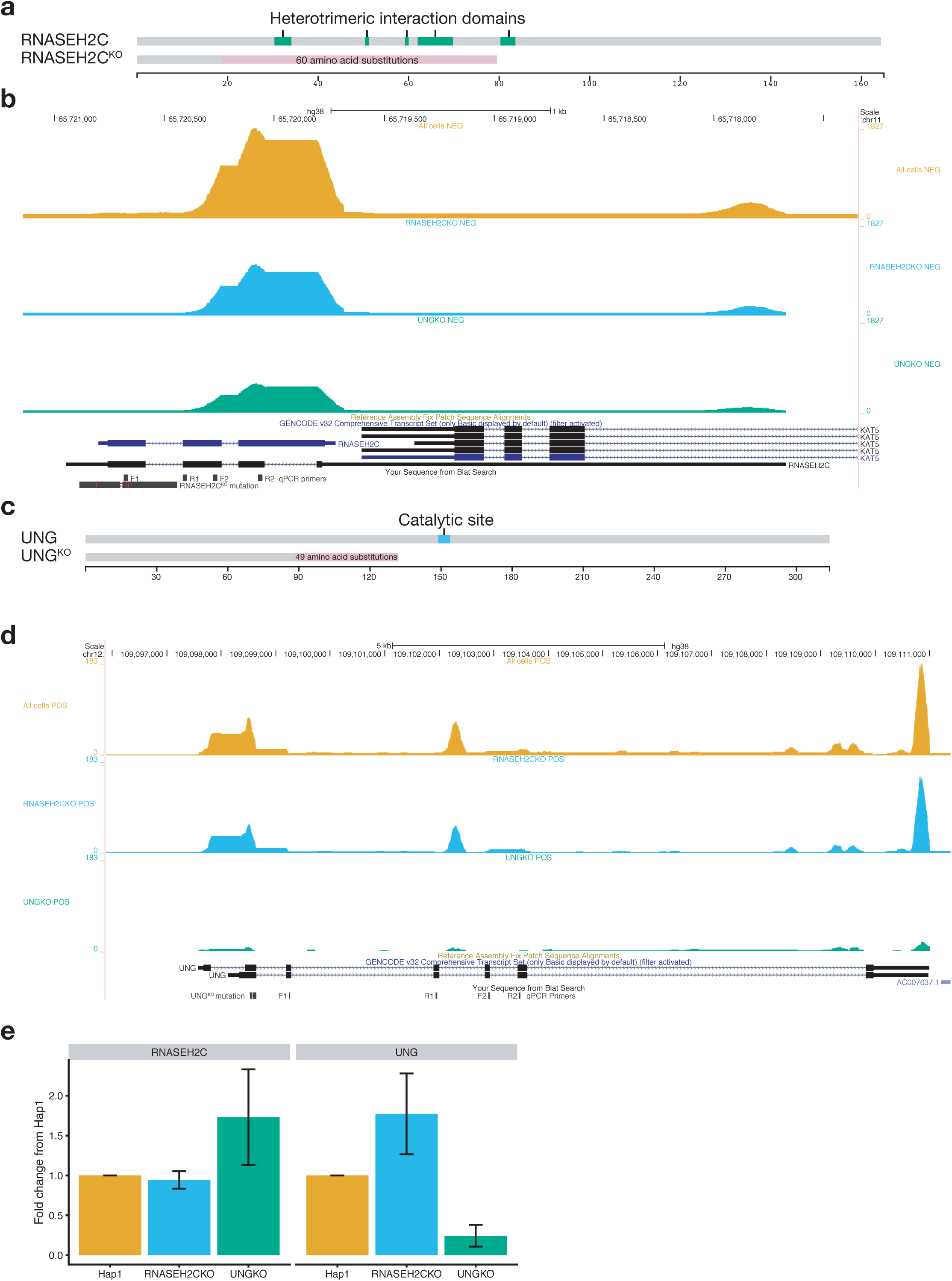
Single-cell mRNA expression of RNASEH2C and UNG in Hap1 knockout cells. **a.** Protein domains for RNASEH2C (top) and mutant RNASEH2C (bottom) from Horizon knockout line. A 10 base pair deletion in the RNASEH2C^KO^ Hap1 line causes a frameshift at amino acid 19 that leads to 60 amino acid substitutions and a stop codon at position 79, disrupting the heterotrimeric interaction regions of RNASEH2C ^25^. **b.** mRNA coverage of RNASEH2C from single-cell RNA sequencing data (**Fig. 1**). Coverage from all cells from negative strand is in orange. Cell types were identified by repair of uracil-containing or ribonucleotide-containing substrates (**Fig. 1**). Alignment files were then separated by cell barcodes associated with cell types. Bulk mRNA coverage for RNASEH2C for RNAESH2C^KO^ cells (blue) and UNG^KO^ cells (green) show similar coverage for the RNASEH2C gene independent of cell type. Coverage for the RNASEH2C gene is located near the 3′ end of the 2 mRNA isoforms (gene diagrams on bottom). Horizon RNASEH2C^KO^ sequencing results provided from Horizon Discovery displayed on bottom (RNASEH2C^KO^ mutation). qPCR primers are also displayed on bottom (F1/2, R1/2). **c.** Protein domains for UNG (top) and mutant UNG (bottom) from Horizon knockout line. An 11 base pair deletion leads to a frameshift at amino acid 88, 49 amino acid substitutions, and a stop codon at amino acid 138 - likely leading to a truncated protein lacking the catalytic domain. **d.** mRNA coverage of UNG from single-cell RNA sequencing data. Coverage from all cells from positive strand is in orange. Cells were identified by repair of uracil-containing or ribonucleotide-containing substrates. Alignment files were then separated by cell type. mRNA coverage for RNASEH2C^KO^ cells (blue) and UNG^KO^ cells (green) show reduced UNG mRNA coverage in UNG^KO^ cells. Gene diagram is on bottom. Horizon UNG^KO^ sequencing results provided from Horizon Discovery is displayed on bottom (UNG^KO^ mutation). qPCR primers are also displayed on bottom (F1/2, R1/2). **e.** Quantitative PCR results for RNASEH2C and UNG mRNA confirm single-cell RNA sequencing results. RNASEH2C^KO^ cells express RNASEH2C mRNA. UNG^KO^ cells do not express UNG mRNA at high levels. Error bars represent errors across biological duplicates and 2 sets of primers indicated in **b** and **d**.

**Supplementary Figure 4.**
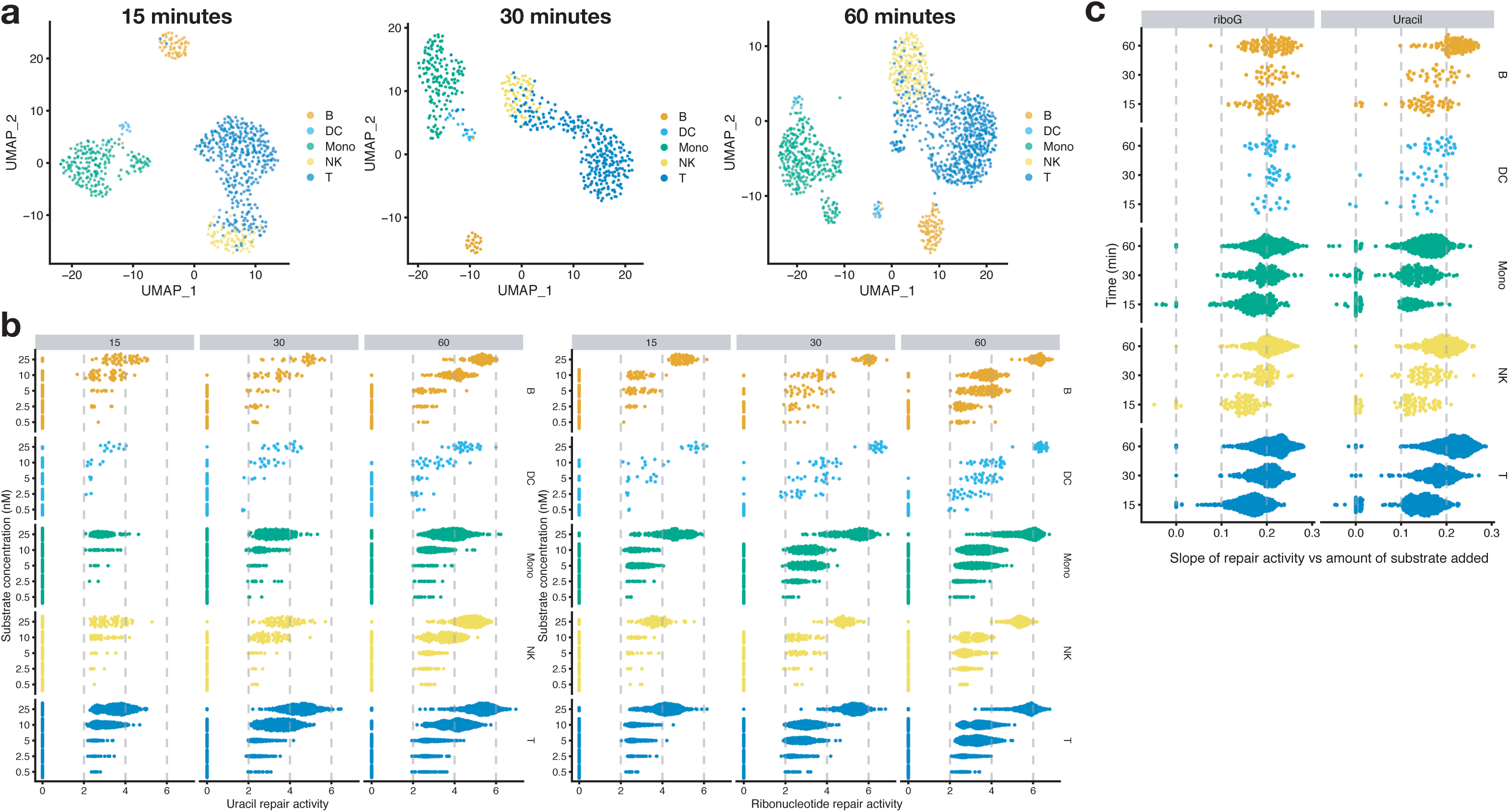
Measuring DNA repair in single cells across multiple concentrations and time points. **a.** DNA repair activities were measured in healthy human PBMCs. Uracil (U:A) and ribonucleotide (rG:C) substrates were added in a range of concentrations (0.5, 2.5, 5, 10, and 25 nM). After the emulsion was created, the sample was separated into 3 tubes and incubated for 15, 30, or 60 min at 37 °C prior to reverse transcription at 53 °C. 800-1,500 cells were captured at each timepoint. Cell types were identified using gene expression markers using Seurat and visualized on UMAP projections. **b.** Repair activities measured increased as substrate concentration increased. Repair activity increased over time from 15 min to 60 minutes. Differences in repair between cell types were consistent across substrate concentrations and time. **c.** A linear model (log normalized count at repair site (as defined in **Fig. 2**) ∼ substrate concentration) was fitted for each cell for each repair activity. The slope of each linear model is a measurement of repair activity. Repair activity increases as time increases. Additionally, repair activity trends measured using a single concentration and time point are consistent with activity measurements made across substrate concentration (e.g., monocytes have lower uracil repair compared to B cells and T cells).

**Supplementary Figure 5.**
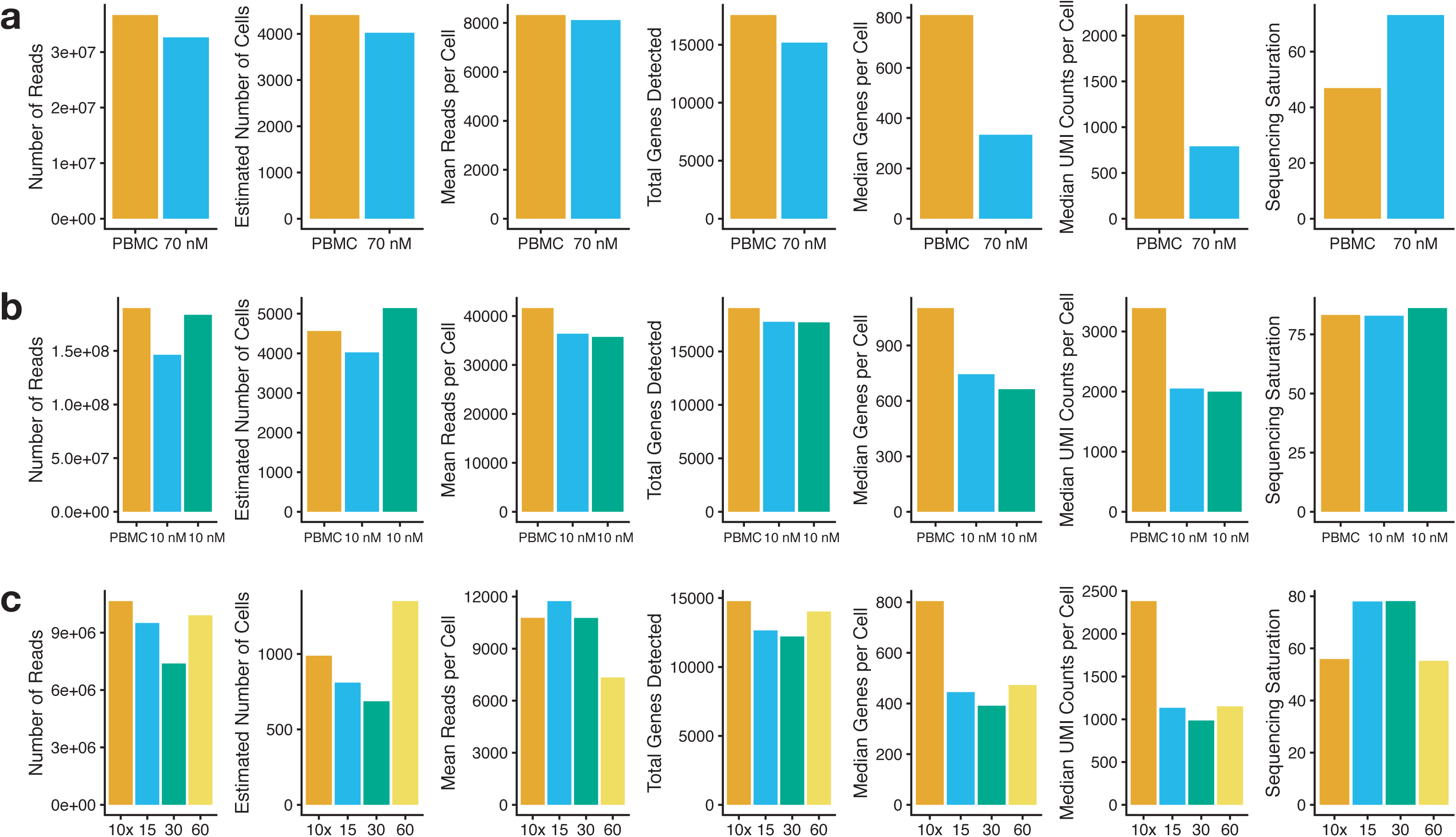
Quality control metrics for single-cell Haircut. **a.** Quality control metrics from 10x cellranger pipeline for PBMC cells with and without the addition of 70 nM DNA repair substrates (**Fig 2**). PBMC data without the addition of repair substrates was downloaded from 10x Genomics v2.1.0 4k PBMC from a healthy human donor (https://support.10xgenomics.com/single-cell-gene-expression/datasets/2.1.0/pbmc4k). Fastq files from 10x Genomics were subsampled to have the same read depth as our PBMC sample (36 million reads). Total genes detected were relatively unaffected by the addition of 70 nM DNA repair substrates, however, median genes per cell and UMI counts per cell was reduced by > 50% with the addition of 70 nM DNA repair substrates. Sequencing saturation was approximately 25% higher in the samples with 70 nM DNA repair substrates. **b.** Quality control metrics from 10x cellranger pipeline for PBMC cells with and without the addition of 10 nM DNA repair substrates in duplicate (**Supplementary Fig 6 and 7**). PBMC data without the addition of repair substrates was downloaded from 10x Genomics v2.1.0 4k PBMC from a healthy human donor (https://support.10xgenomics.com/single-cell-gene-expression/datasets/2.1.0/pbmc4k). Fastq files from 10x Genomics were subsampled to have the same read depth as our PBMC sample (160 million reads). Median genes per cell and UMI counts per cell was reduced by ∼40% with the addition of 10 nM DNA repair substrates. **c.** Quality control metrics from 10x cellranger pipeline for PBMC cells with and without the addition of 110 nM DNA repair substrates (**Supplementary Fig 4**). Samples containing DNA repair substrates were incubated at 37 °C for 15, 30, or 60 minutes prior to incubation at 53 °C for reverse transcription. PBMC data without the addition of repair substrates was downloaded from 10x Genomics v3.0.0 1k PBMC from a healthy human donor v2 (support.10xgenomics.com/single-cell-gene-expression/datasets/3.0.0/pbmc_1k_v2). Fastq files from 10x Genomics were subsampled to have the same read depth as our PBMC sample (10 million reads). Median genes detected per cell and UMI counts per cell was reduced by ∼50% with the addition of 110 nM DNA repair substrates. This decrease in genes detected and UMI counts was independent of incubation time at 37 °C.

**Supplementary Figure 6.**
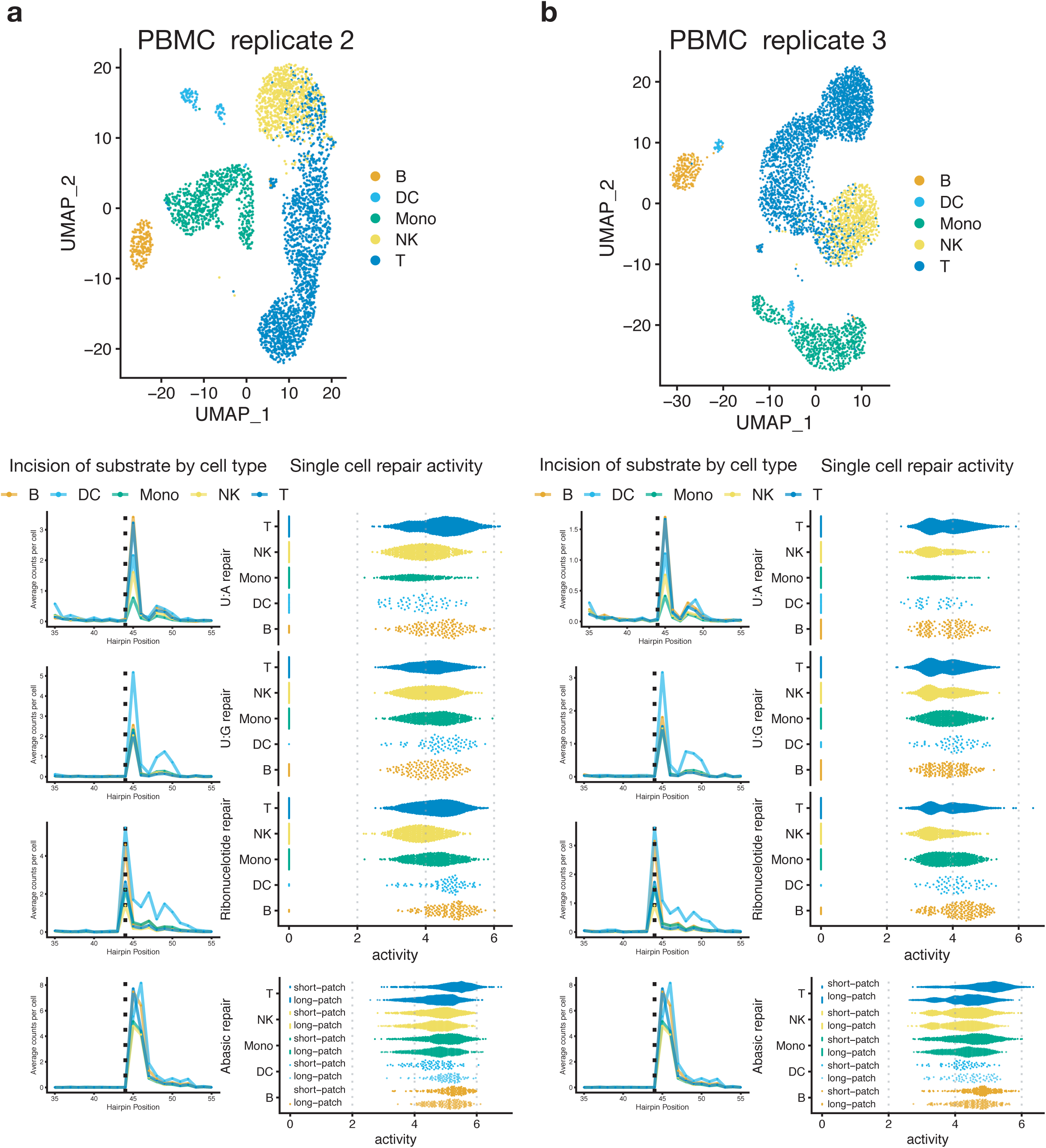
Biological replication of DNA repair phenotypes in human PBMCs. **a-b.** PBMCs were isolated from a single healthy human donor in two batches. 3′ Single cell gene expression and DNA repair activities were measured for each batch. mRNA expression was used to cluster cells and classify cell types. UMAP plot of cell types (top). Cell-type-specific counts of incision and processing (mean) are plotted against the position of the hairpin (left). Trends for single cell repair of U:A, U:G, riboG:C, and abasic:G substrates were similar across all three replicates (**Fig. 2 and a-b**).

**Supplementary Figure 7.**
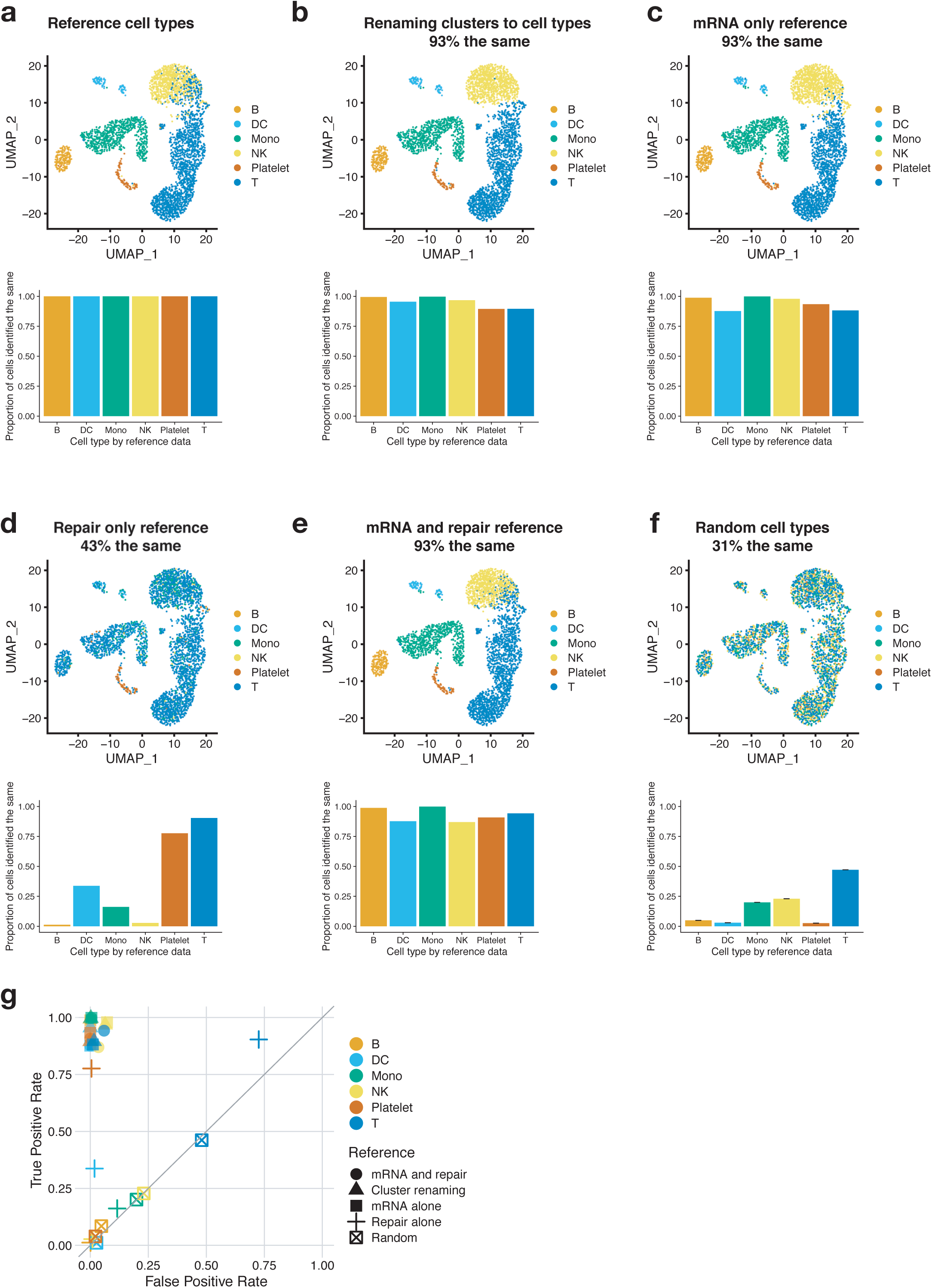
Cell-type classification using DNA repair measurements. **a.** PBMC UMAP and cell classifications for PBMC2 replicate (**Supplementary Fig. 6**). Cells were classified using reference data from 10x Genomics and Seurat v3.0.0 FindTransferAnchors and TransferData functions. These cell classifications are used as the true cell type for other classification methods, however, these classifications may not be 100% accurate. The percentage of each cell type that is classified the same as this reference data (bottom) is 100% since it is compared to itself. **b.** Seurat clusters (0-10) were renamed for the majority cell type marker from **a** in each cluster. UMAP plot of renamed cells (top). Using this classification method 93% of cells are classified the same as **a**. > 89% of each cell type were classified the same (bottom). **c.** mRNA data in **Fig. 2** was used as the reference for classifying cell types in PBMC2 replicate (**Supplementary Fig. 6**) using Seurat as before (**a**). Using this reference, 93% of the cells were classified the same. > 87% of each cell type were classified the same (bottom). **d.** DNA repair measurements alone from PBMCs (**Fig. 2**) was used as the reference for classifying cell types PBMC2 replicate. Using only DNA repair for reference data, only 43% of cells were classified the same as **a**. Platelets lack DNA repair activities in Haircurt (data not shown) which could contribute to their identification using DNA repair measurements alone as they are distinctly different from all other cells. > 90% of T cells were also identified using repair alone, however, T cells make up ∼40% of all cells in the data and when classified using repair data alone, ∼80% of all cells are classified as T cells, so by chance alone,T cells are more likely to be classified correctly. Dendritic cells make up only 2.6% of all cells in the data, however, 33% of them are classified correctly using repair activity data alone. DC have several unique repair signatures (**Fig. 2** and **Supplementary Fig. 6**) which are likely to contribute to their classification using DNA repair activities alone. **e.** Count matrices for mRNA expression and DNA repair were combined and used as a reference (from **Fig. 2** data) to classify PBMC2 replicate cells. Using this reference, 93% of the cells were classified the same as in **a**. > 86% of each cell type were classified the same (bottom). **f.** Cell type labels from PBMC2 as defined in **Supplementary Fig. 6** were randomly reassigned to cell ids. Only 31% of cells were classified the same. More abundant cell types were more likely to be correctly classified by chance (bottom). Error bars represent the 95% confidence interval for 1000 independent samplings. **g.** True positive rate (# of true positives / (# of true positives + # of false negatives)) and false positive rate (# of false positives / (# of false positives + # of true negatives)) for each cell type classified by each method. Cells classified using mRNA expression with or without repair had a high true positive rate and low false positive rate. Most cell types defined by repair alone had relatively equal true and false positive rates, with the exception of platelets which had a high true positive and low false positive rate and DC which had a relatively low true positive rate but a very low false positive rate indicating DNA repair measurements for these cell types may be helpful for classification.

**Supplementary Figure 8.**
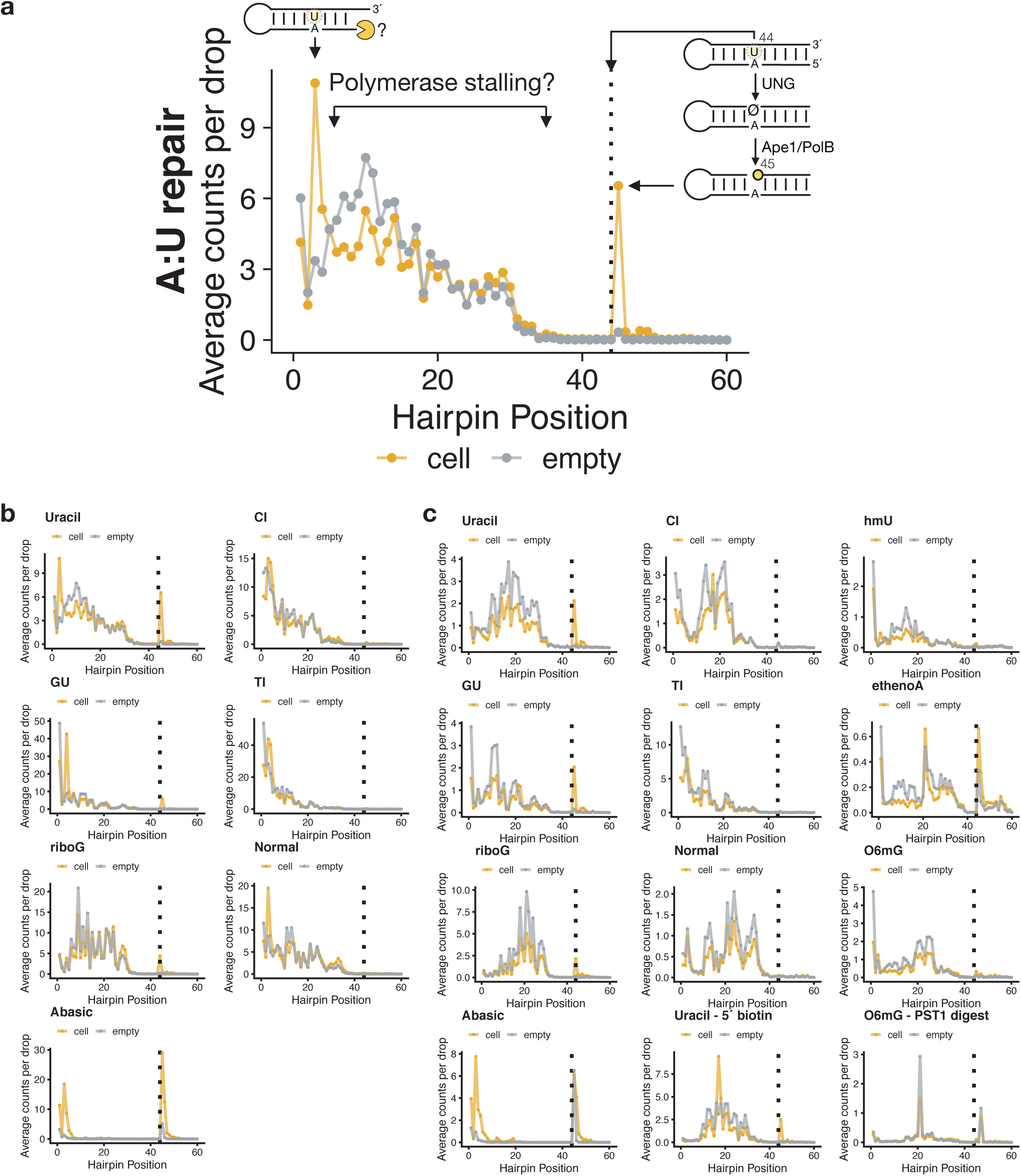
Determining biological DNA repair activity from single-cell Haircut signals. **a.** Counts per cell (mean) (orange) by position were compared to counts per empty drop (mean) to determine which positions contained biological activity. Empty drops were determined by filtering out cell-associated barcodes from the unfiltered repair matrix. The resulting repair matrix contains many barcodes that are associated with only a single UMI, so the matrix was filtered by descending UMI counts to the same number of cell-associated barcodes. This repair matrix was used as the input to calculate empty drop signal across the hairpin by calculating the sum across all drops at each hairpin position. Positions exhibiting signal above empty drop background and associated with a known DNA repair position (e.g., position 45 for U:A) were considered biologically relevant. Some signal above background at the 5′ end of the substrate could be due to cellular exonuclease activity and were not considered in the analysis. **b.** Coverage across all cells (orange) and empty drops (grey) for PBMC experiment (**Fig. 2**). C:I and T:I substrates did not show biological activity above background. **c.** Coverage across all cells (orange) and empty drops (grey) for PBMC experiments (**Supplementary Fig. 6a**). The Uracil-5′ biotin substrate contained a 5′-biotin. During the library preparation, uncleaved substrate was removed using streptavidin beads. We only saw a modest reduction in signal on the 5′ end of the substrate in cell-containing and empty drops. The hmU substrate did not have signal above empty drop background. The ethenoA substrate did not have signal above background due to a high level of background signal in empty drops, which could be due to the bulky ethenoA lesion causing the RT to stop. To measure direct reversal substrates, we included an O6mG containing hairpin. Following substrate isolation from the mRNA fraction, substrates were digested with PstI to measure the removal of O6mG ^26^; however, we measured digestion in empty drops as well as drops with cells indicating the method was not specific for droplets containing cells.

